# High-Quality Genomes of Nanopore Sequencing by Homologous Polishing

**DOI:** 10.1101/2020.09.19.304949

**Authors:** Yao-Ting Huang, Po-Yu Liu, Pei-Wen Shih

## Abstract

Nanopore sequencing has been widely used for reconstruction of a variety of microbial genomes. Owing to the higher error rate, the assembled genome requires further error correction. Existing methods erase many of these errors via deep neural network trained from Nanopore reads. However, quite a few systematic errors are still left on the genome. This paper proposed a new model trained from homologous sequences extracted from closely-related genomes, which provides valuable features missed in Nanopore reads. The developed program (called Homopolish) outperforms the state-of-the-art Racon/Medaka and MarginPolish/HELEN pipelines in metagenomic and isolates of bacteria, viruses and fungi. When Homopolish is combined with Medaka or with HELEN, the genomes quality can exceed Q50 on R9.4 flowcells. The genome quality can be also improved on R10.3 flowcells (Q50-Q90). We proved that Nanopore-only sequencing can now produce high-quality genomes without the need of Illumina hybrid sequencing.

## Background

The third-generation long-read sequencing is an essential technology for reconstruction of complete genomes of many species in the biosphere. The Oxford Nanopore Technology (ONT) is one of the major providers in third-generation sequencing, which is being used for reconstructing human chromosomes from telomere to telomere [1, 2]. Although the ultra-long reads of ONT has demonstrated its power in assembly contiguity through large and complex repeat regions in the genome, its assembly accuracy (∼85-92%) has been criticized in comparison with Illumina or PacBio High-Fidelity (HiFi) sequencing (∼99%), owing to miss of important protein-coding genes [3]. As a consequence, hybrid Illumina/Nanopore sequencing and assembly is often performed in order to produce a high-quality genome having both high contiguity and accuracy [4].

The accuracy of ONT reads improved year by year thanks to new basecalling algorithms (e.g., Albacore to Guppy or to Bonito) and flowcells (e.g., R9.4 to R10.3). For instance, their production basecaller (Guppy v3.6) claimed 1-2% increase in accuracy (∼97%) than the previous version. However, the genomes assembled from ONT raw reads are still far from satisfactory accuracy owing to a substantial number of systematic errors. Consequently, all the long-read assemblers (e.g., Canu, miniasm, Flye, Shasta) require a polishing stage for further improving the genome quality [5, 6, 7, 8]. At the beginning, signal-based polishing (e.g., Nanopolish) were adopted, whereas potential erroneous loci were re-basecalled from raw signals [9]. But it is time-consuming and disk-demanding for processing and for storing the huge amount of signals. Nowadays, read-based polishing becomes the mainstream as only reads instead of signals are required. Thanks to the advances of combinatorial and machine-learning algorithms, they are not only faster but also produces higher accuracy than signal-based approaches. Below we focused on two state-of-the-art read-based polishing pipelines: Racon/Medaka and NarginPolish/HELEN.

Racon is one of the most popular polishing programs [10]. It first carefully selects high-quality parts of reads, and then the genome is polished via Partial Order Alignment (POA) with vectorization. Although many errors are corrected by Racon, a substantial amount of systematic errors is left on the genome, because the correct allele is a minority at these loci (Supplementary Figure S1). As a consequence, the Oxford Nanopore Inc. developed Medaka, which is based on a bidirectional Long-Short-Term Memory (LSTM) trained for erasing systematic errors missed by Racon. To date, polishing first by Racon and followed by Medaka is the officially recommended protocol (i.e., Pomoxis) for genomes assembled solely from Nanopore sequencing.

Recently, a new polishing pipeline called MarginPolish and HELEN drew attention by showing competitive accuracy compared with the Racon/Medaka pipeline [8]. MarginPolish uses a hidden Markov model to collect alignment statistics and generates a weighted POA graph for consumption by HELEN. Subsequently, HELEN uses a multi-task recurrent neural network (RNN) that utilizes both the contextual genomic features and POA weights to predict a nucleotide base and run length for each genomic position with high accuracy.

However, these polishing protocols still failed to guarantee a high-quality genome can be produced. In reality, (∼Q30) (99.9%) consensus accuracy can be obtained, which can still miss important genes in downstream analysis [11, 3]. The unsatisfactory quality is partly due to species-specific DNA modifications. Theoretically, these errors might be further removed by training a basecaller specific for each species (e.g., Taiyaki) [11]. But practically, it is infeasible to train thousands of basecalling models for all the species in the biosphere. We observed homologous sequences from closely-related genomes provide valuable features missed in Nanopore reads and all previous works (Supplementary Figure S2). For instance, indels within coding regions are extremely rare as they lead to catastrophic frameshift. Consequently, existing polishing models can be easily improved by capturing the conservative sequence context within homologous regions.

This paper developed a novel polishing tool (called Homopolish) based on a support-vector machine trained from homologous sequences extracted from closely-related genomes. With carefully engineered features, the results indicate that Homopolish outperforms state-of-the-art Medaka and HELEN pipelines over a variety of public and in-house sequenced genomes, including metagenomic, bacterial, viral and fungal genomes.

## Results

Homopolish is a machine learning model trained for correcting Nanopore systematic errors, indel errors in particular, by exploring sequence conservation context across homologous sequences. We compare Homopolish with two state-of-the-art polishing pipelines (Racon/Medaka and MarginPolish/HELEN) [8, 10]. These programs were evaluated using public or in-house sequenced metagenomic/isolate datasets, including bacteria, virus, and fungi (Supplementary Tables S1-S3, see Methods). The Nanopore reads of all datasets were first assembled into draft genomes by Flye or MetaFlye (Supplementary Tables S4-S6) [7, 12]. The draft genomes were first corrected by Racon or by MarginPolish for removing random errors, and subsequently polished by Medaka, by HELEN, and by Homopolish for removing systematic errors. The accuracy of the polished genome is measured by (median) Q score, number of mismatches, number of insertions, and number of deletions using fastmer [13].

### Comparison of genome quality on R9.4 metagenomic datasets

We first compare Medaka over Racon, Homopolish (R) over Racon, HELEN over MarginPolish, and Homopolish (M) over MarginPolish on a metagenomic dataset (Zymo Microbial Community standard) sequenced by R9.4 flowcells. Figure 1 lists the Q scores of all programs over seven bacteria within the metagenomic sample, where the numbers of insertion, deletions, and mismatches can be found in Supplementary Table S7. Homopolish (R) and Homopolish (M) (∼Q38-Q50) outperforms Medaka (∼Q36-38) and HELEN (∼Q37-Q46) in most bacteria. Homopolish (M) achieves Q50 (99.999%) in *Enterococcus faecalis, Pseudomonas aeruginosa*, and *Salmonella enterica*. The only exception is *B subtilis* (Q36.78 for Homopolish and Q37.21 for Medaka), which is due to mismatches which are not corrected by Homopolish, although indels are greatly reduced (Supplementary Figure S3). In general, systematic errors are greatly reduced by Medaka, Helen, and Homopolish in comparison with those solely by Racon and by MarginPolish. Nevertheless, we observed results based on MarginPolish (i.e., HELEN and Homopolish(M)) are superior to those based on Racon (i.e., Medaka and Homopolish (R)).

**Figure 1.**
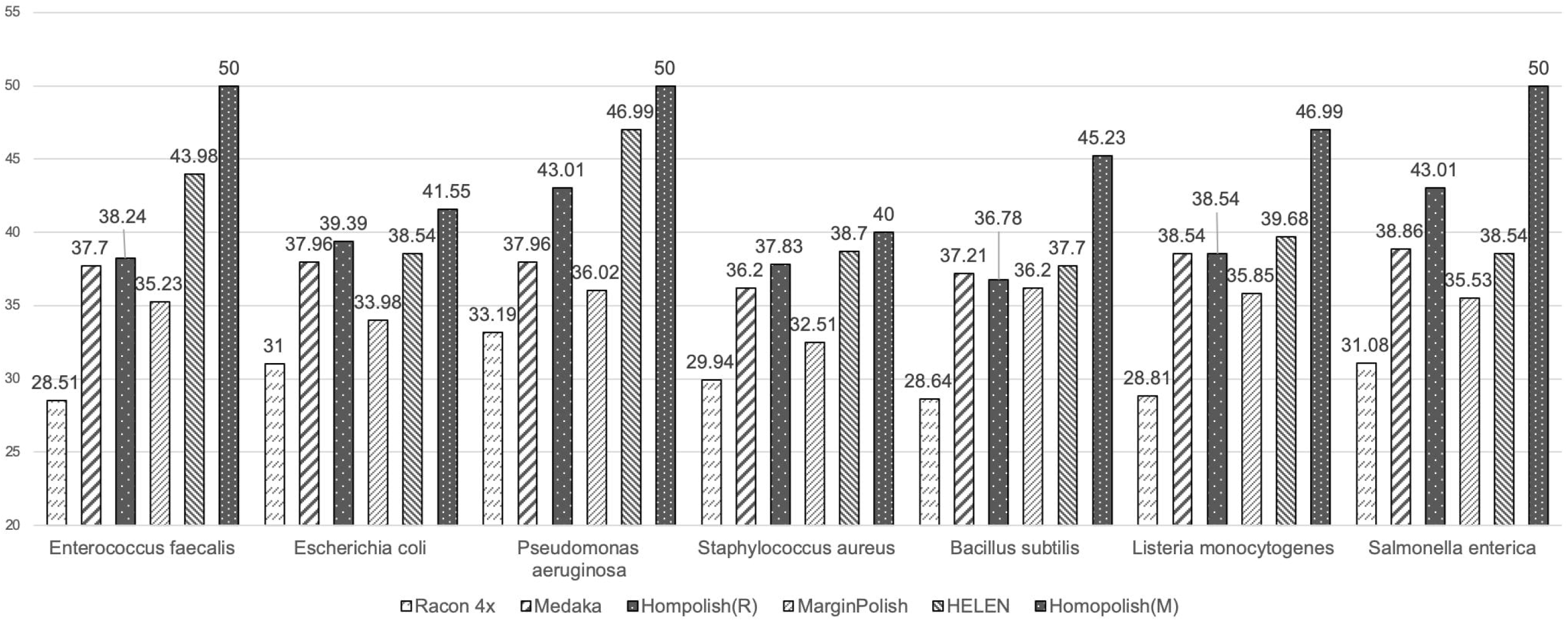
Comparison of genome quality on the R9.4 metagenomic dataset. Comparison of genome quality (Q score) polished by Racon, Medaka, Homopolish(R), MarginPolish, HELEN, and Homopolish(M) on the metagenomic dataset from Zymo Microbial Community Standard. Medaka and Homopolish(R) are run after Racon. HELEN and Homopolish(M) are invoked after MarginPolish. Homopolish (M) and Homopolish (R) achieve highest accuracy in most bacteria.

Because mismatched errors were not corrected by Homopolsh, we want to know if Homopolish can achieve even higher accuracy when combining with Medaka or HELEN. Figure 2 plots the Q scores of invoking Homopolish after Medaka and HELEN polishing. The accuracy of Medaka and HELEN are both further improved by Homopolish. For instance, Homopolish after Medaka now reaches Q50 on *Salmonella enterica* and exceeds Q40 for most bacteria. In general, Homopolish with HELEN (Q41-Q90) outperforms that with Medaka (Q39-Q50). In particular, Homopolish after HELEN achieves Q90 on *Pseudomonas aeruginosa* and Q50 on *Enterococcus faecalis* and on *Salmonella enterica*.

**Figure 2.**
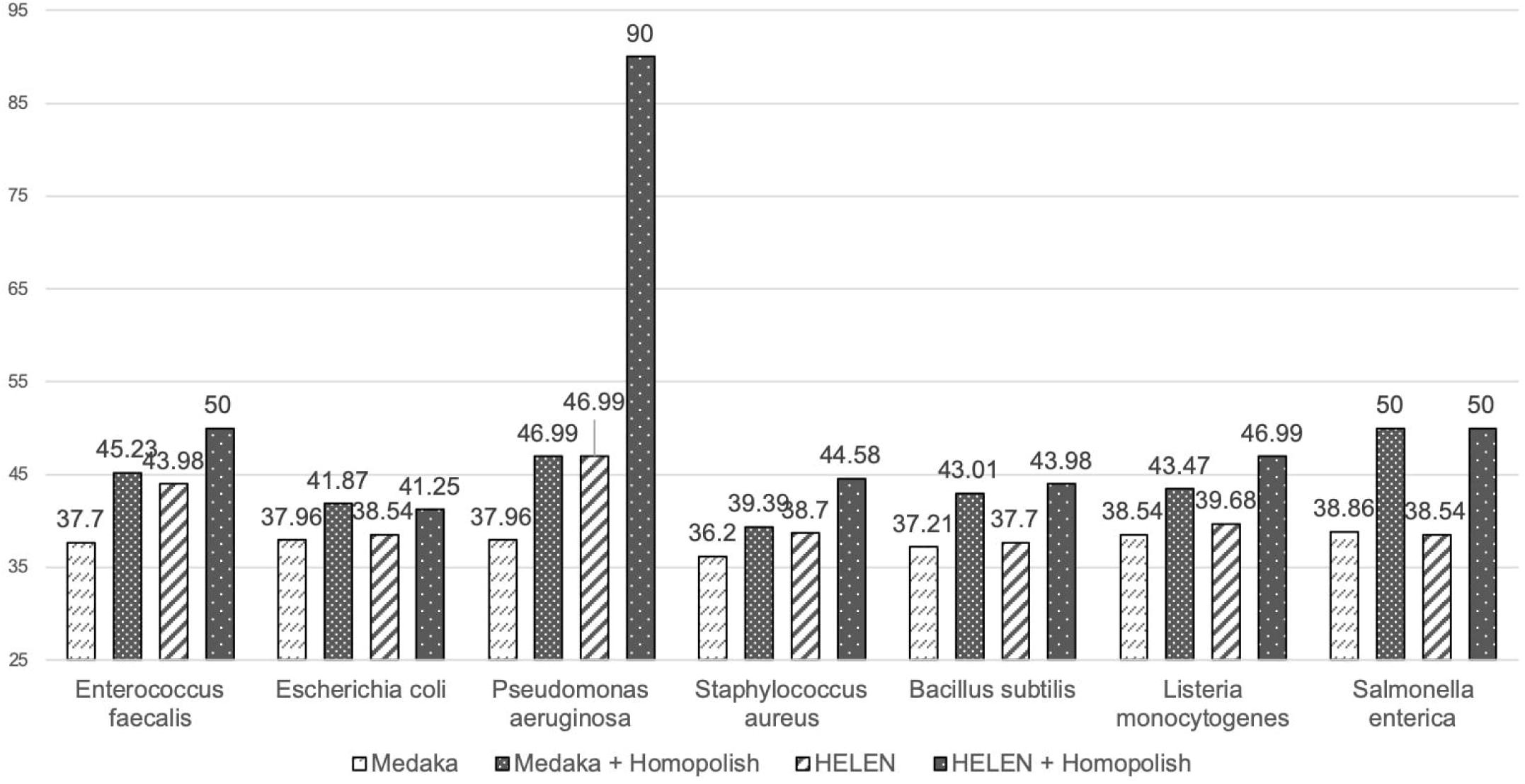
Genome quality of combining Homopolish with Medaka or with HELEN in the metagenomic dataset. Comparison of genome quality (Q score) polished by Medaka, Medaka+Homopolish, HELEN, and HELEN+Homopolish on the metagenomic dataset. Medaka and HELEN can be both further improved by Homopolish.

Table 1 lists the numbers of mismatches, insertions, and deletions of the *Pseudomonas aeruginosa* genome polished by all programs, where those of other six bacteria can be found in Supplementary Table S7. The quality of draft genome (assembled and polished by Flye) is only around Q26, and the errors are dominated by 14,394 insertions. Racon removed quite a few insertion errors at the cost of producing many false deletion errors (from 439 to 2,417). Medaka corrected most of deletion errors produced by Racon (from 2,417 to 324) and a few mismatches (from 622 to 476). Homopolish significantly erased most insertion and deletion errors (from 538 and 2,417 to 24 and 128, respectively) left by Racon. These phenomena are largely the same when Homopolish was compared with HELEN over MarginPolish. We found HELEN is superior at correcting mismatched errors and outperforms Medaka at all metrics. Again, Homopolish cleans the majority of indel errors left by MarginPolish. Consequently, Homopolish still outperforms HELEN and Medaka in most datasets (Supplementary Table S7). When Homopolish is combined with Medaka or with HELEN, the genome quality is further elevated to Q47 or to Q90, respectively. The is because both mismatches and indel errors are removed significantly by each program.

**Table 1.**
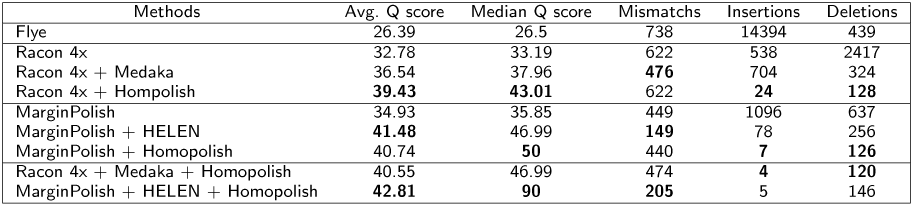
Q scores and numbers of mismatch, insertion, and deletion errors of *P aeruginosa* in the metagenomic dataset.

| Methods | Avg. Q score | Median Q score | Mismatches | Insertions | Deletions |
| --- | --- | --- | --- | --- | --- |
| Flye | 26.39 | 26.5 | 738 | 14394 | 439 |
| Racon 4x | 32.78 | 33.19 | 622 | 538 | 2417 |
| Racon 4x + Medaka | 36.54 | 37.96 | **476** | 704 | 324 |
| Racon 4x + Homopolish | **39.43** | **43.01** | 622 | **24** | **128** |
| MarginPolish | 34.93 | 35.85 | 449 | 1096 | 637 |
| MarginPolish + HELEN | **41.48** | 46.99 | **149** | 78 | 256 |
| MarginPolish + Homopolish | 40.74 | **50** | 440 | **7** | **126** |
| Racon 4x + Medaka + Homopolish | 40.55 | 46.99 | 474 | **4** | **120** |
| MarginPolish + HELEN + Homopolish | **42.81** | **90** | **205** | 5 | 146 |

### Comparison of polishing accuracy on bacterial isolates

Next, We compare these programs over a set of bacterial isolates sequenced at earlier stage (see Methods). They were mainly obtained before 2018 using Albacore and/or early Guppy (2.x) basecaller, which exhibited lower quality compared with previous metagenomic dataset (called by Guppy 3.x). Theoretically, old sequencing data can be re-called using new basecalling algorithms. But practically, especially for labs outsourcing the entire sequencing and assembly service, this is very troublesome as the assembled genomes are the major material. We show that Homopolish can improve the quality of early-generation genomes without the need of even signals or reads. Figure 3 plots the Q scores of all programs across seven isolates. Similarly, Homopolish (Q26-Q33) outperforms Medaka (Q23-Q29) over Raon in nearly all datasets, except for *K penumoiae* in which mismatches (not corrected by Homopolish) are the major error source (Supplementary Figure S4). When comparison with HELEN over MarginPolish (Q20-Q27), Homopolish (M) also showed superior accuracy (Q26-Q37) across all isolates. Unexpectedly, the accuracy of HELEN is not only lower than Medaka but also than its preprocessor MarginPolish in most datasets (i.e., *E coli, K pneumonia, E anopheies* and *S algae*), which is in conflict with previous metagenomic results. We hypothesize HELEN may overfit previous metagenomic dataset as majority of its training data come from the same source (ZymoBIOMICS) [14].

**Figure 3.**
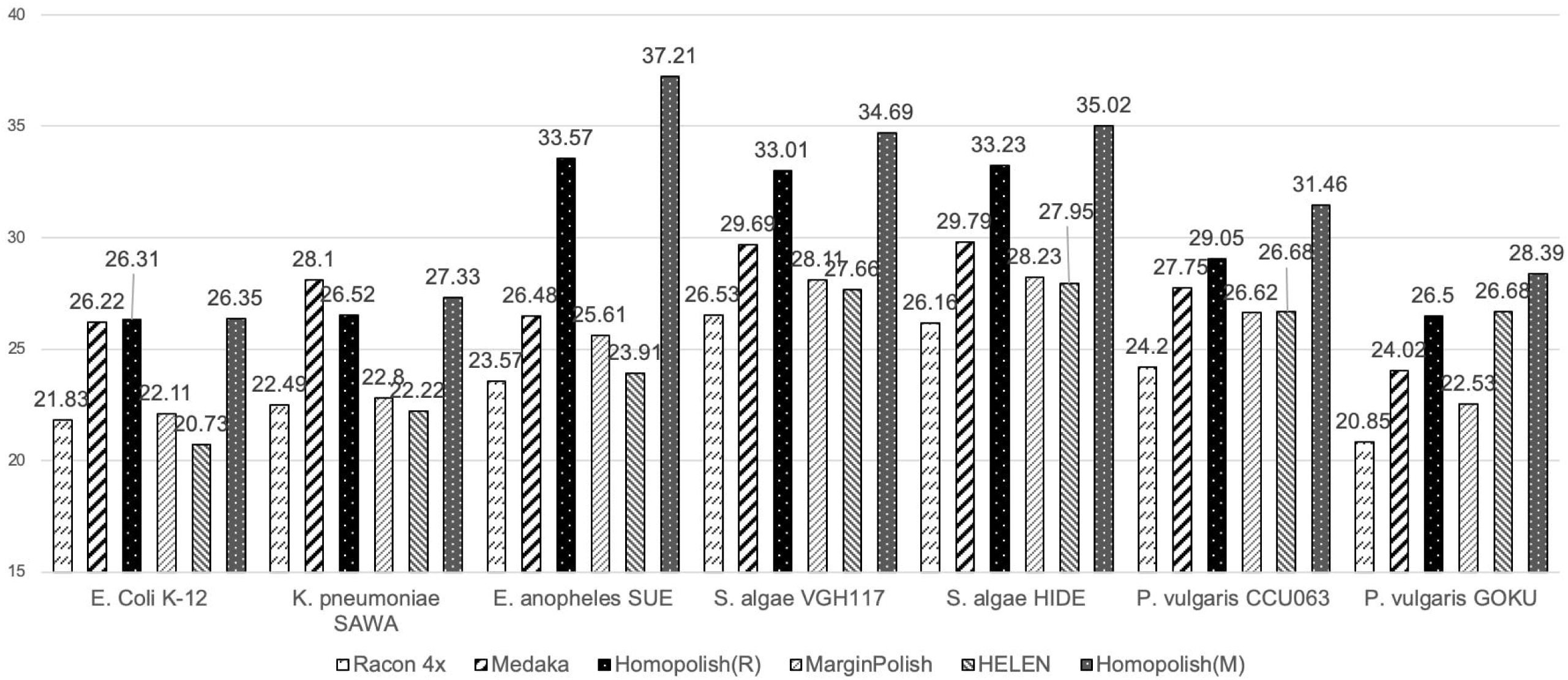
Comparison of genome quality on the R9.4 isolate dataset. Comparison of genome quality (Q score) polished by Racon, Medaka, Homopolish(R), MarginPolish, HELEN, and Homopolish(M) on seven bacterial isolates. Medaka and Homopolish(R) are run after Racon. HELEN and Homopolish(M) are invoked after MarginPolish.

Similarly, we evaluate whether Homopolish can achieve even higher accuracy when combined with Medaka or with HELEN over these isolates. Figure 4 illustrates the Q scores of Homopolish run after Medaka or HELEN correction. When combined with Medaka, Homopolish obtains much higher accuracy (Q29-Q38), significantly better than those of original Medaka (Q23-Q29). When Homopolish was run after HELEN, it indeed further improves accuracy from Q20-Q27 to Q24-Q35. However, as HELEN has inferior accuracy on these isolates, Homopolish with Medaka achieves the highest accuracy in these isolates. Table 2 lists the Q scores, number of mismatches, insertions, and deletions polished by each program of *E*.*anopheles SUE*, where those of other isolates can be found in Supplementary Table S8. The initial genome quality of Flye is low, containing 18,718 insertions and 5,039 deletions.

**Table 2.**
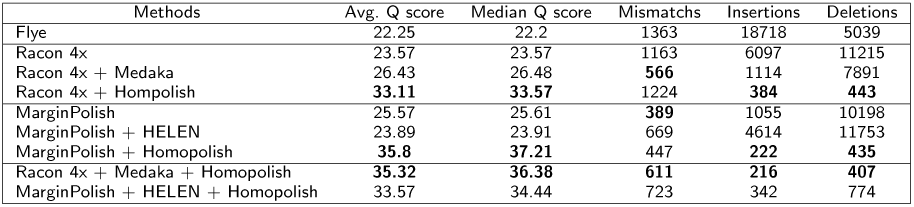
Q scores and numbers of mismatch, insertion, and deletion errors of *E*.*anopheles SUE* in isolates datasets

**Figure 4.**
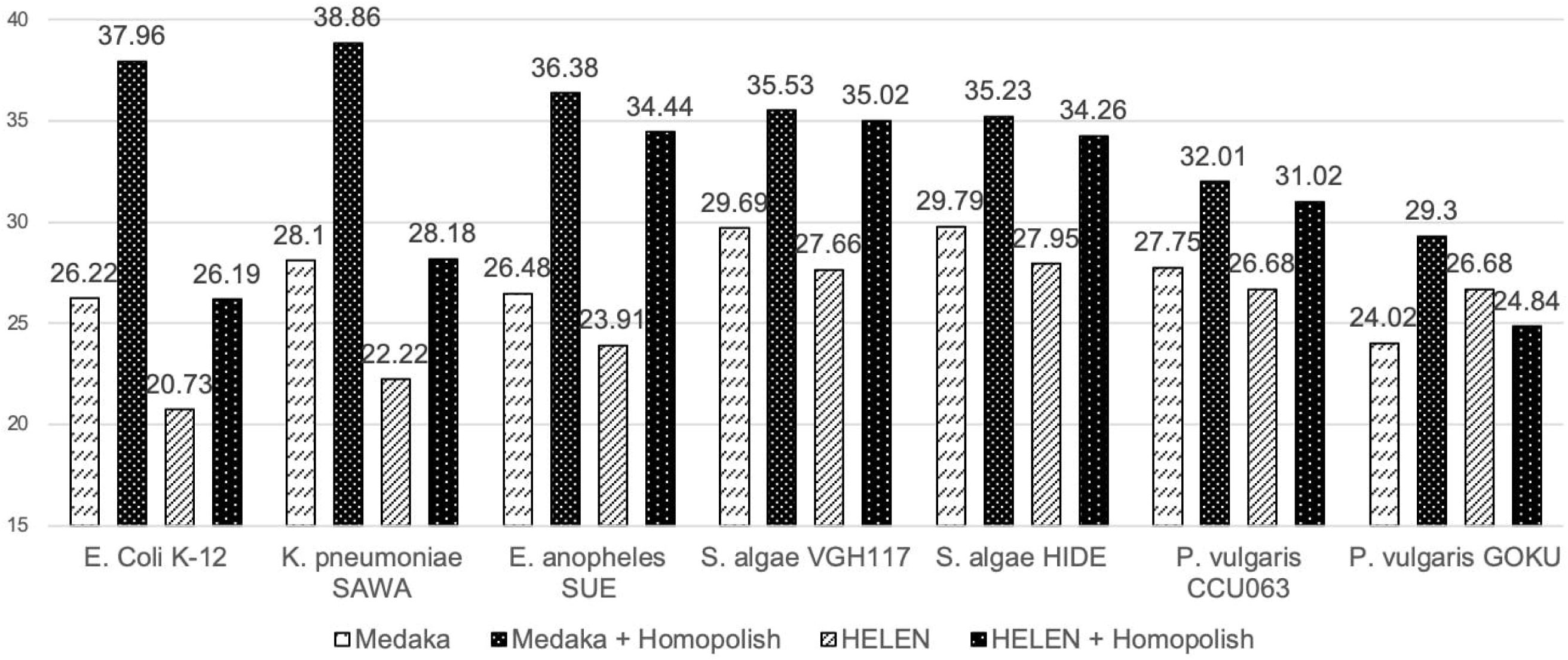
Genome quality of combining Homopolish with Medaka or with HELEN in the isolate dataset. Comparison of genome quality (Q score) polished by Medaka, Medaka+Homopolish, HELEN, and HELEN+Homopolish on the isolate dataset. Medaka and HELEN can be both further improved by Homopolish.

Although Racon, Medaka, MarginPolis, and HELEN reduced the insertion errors, the numbers of deletion errors are all increased (i.e., from 5,039 to 7,891-11,753), implying quite a few false-positive corrections. On the contrary, only Homopolish didn’t introduce more false deletions into the draft genome (i.e., from 5,039 to 435-443). In particular, HELEN performs worst due to increasing indel errors from its preprocessor MarginPolish in most isolates (Supplementary Table S8). Overall, only Homopolish can correct most indel errors without much side effects (222-384 insertions and 435-443 deletions). When combined with Medaka for removing both mismatched and indel errors, Homopolish achieves better accuracy as expected (from Q33 to Q36). But the combination of Homopolish with HELEN (Q34) was not better than running Homopolish directly on top of MarginPolish (Q37).

### Comparison of correction accuracy on viral and fungal genomes

In addition to bacteria, these programs were further tested on one viral (*Lambda phage*) and on one fungal (*S. cerevisiae*) genomes. As shown in Table 3, the draft viral genome is around Q24. Racon with Medaka or with Homopolish both improved the quality. Homopolish achieves the highest accuracy when combined with Racon or with Medaka (from ∼Q24 to ∼Q38-39). Accuracy of HELEN is not only worst but also lower than original draft genome (from ∼Q24 to ∼Q20), owing to numerous mismatches and deletion errors falsely generated. In *S. cerevisiae* fungal genome, the quality of initial genome is about Q20. Correction via Racon and Medaka improved the quality to Q23. Highest accuracy were achieved by Homopolish when combined with Medaka or with MarginPolish (∼Q32). Again, HELEN obtained lowest accuracy in comparison with others. These results indicated that Racon, Medaka, and Homopolish are robust to polished viral, bacterial, and fungal genomes while HELEN is less reliable compared with others.

**Table 3.**
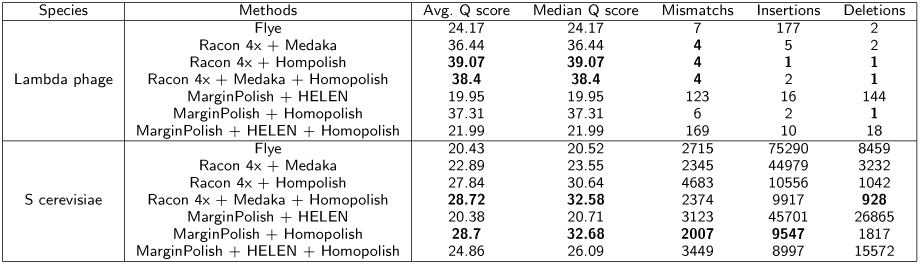
Q scores and numbers of mismatch, insertion, and deletion errors on *Lambda phage* and *Saccharomyces cerevisiae*

| Species | Methods | Avg. Q score | Median Q score | Mismatches | Insertions | Deletions |
| --- | --- | --- | --- | --- | --- | --- |
| Lambda phage | Flye | 24.17 | 24.17 | 7 | 177 | 2 |
|  | Racon 4x + Medaka | 36.44 | 36.44 | **4** | 5 | 2 |
|  | Racon 4x + Homopolish | **39.07** | **39.07** | **4** | **1** | **1** |
|  | Racon 4x + Medaka + Homopolish | **38.4** | **38.4** | **4** | 2 | **1** |
|  | MarginPolish + HELEN | 19.95 | 19.95 | 123 | 16 | 144 |
|  | MarginPolish + Homopolish | 37.31 | 37.31 | 6 | 2 | **1** |
|  | MarginPolish + HELEN + Homopolish | 21.99 | 21.99 | 169 | 10 | 18 |
| S. cerevisiae | Flye | 20.43 | 20.52 | 2715 | 75290 | 8459 |
|  | Racon 4x + Medaka | 22.89 | 23.55 | 2345 | 44979 | 3232 |
|  | Racon 4x + Homopolish | 27.84 | 30.64 | 4683 | 10556 | 1042 |
|  | Racon 4x + Medaka + Homopolish | **28.72** | **32.58** | 2374 | 9917 | **928** |
|  | MarginPolish + HELEN | 20.38 | 20.71 | 3123 | 45701 | 26865 |
|  | MarginPolish + Homopolish | **28.7** | **32.68** | **2007** | **9547** | 1817 |
|  | MarginPolish + HELEN + Homopolish | 24.86 | 26.09 | 3449 | 8997 | 15572 |

### Comparison of genome quality on R10.3 flowcells

Finally, we evaluate Racon, Medaka, and Homopolish on one public metagenomic dataset (Zymo Microbial Community Standard) and one public isolate (E coli K12 MG1655) sequenced by R10.3 flowcells (see Methods). R10.3 is expected to exhibit higher accuracy than R9.4 flowcells at the cost of reduced throughput. HELEN was excluded because of previous overfitting issue. Figure 5 plots the Q scores of Racon, followed by Medaka, and finally by Homopolish on the metagenomic dataset sequenced by R10.3 flowcells. Supplementary Table S9 lists the numbers of mismatches, insertions, and deletions of these bacteria. The quality of draft genomes ranges from Q28 to Q42, which is better than that of R9.4 (Q22-Q26). Unexpectedly, the genomes polished by Racon did not show improvement (Q27-Q40) compared with the draft genome by Flye. Genomes corrected by Medaka showed clear higher quality (from Q30 to Q50). Homopolish further improved the genome from Q33 to Q90 (e.g., Q90 on *E faecalis* and *P aeruginosa*). On the E coli isolate R10.3 dataset (Supplementary Table S9), Homopolish improved Medaka from Q46 to Q90. We note that the accuracy calculation includes not only uncorrected errors but also assembly errors and strain variations. Consequently, the accuracy of genomes sequenced by R9.4 and by 10.3 flowcells does not differ too much (i.e., both can exceed Q50) if polished by Homopolish.

**Figure 5.**
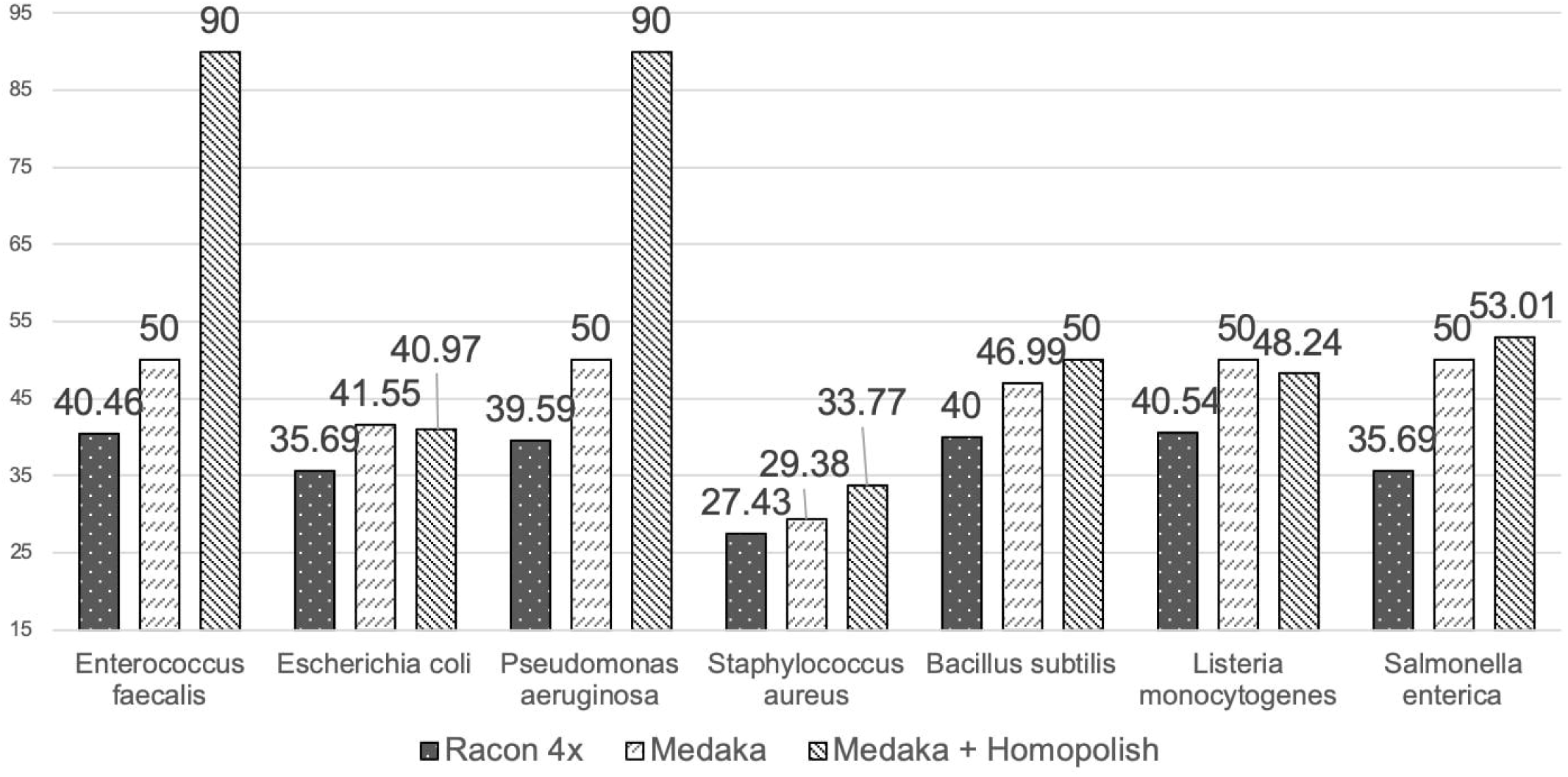
Comparison of genome quality on the R10.3 metagenomic dataset. Comparison of genome quality (Q score) polished by Racon, Medaka, and Homopolish on the R10.3 metagenomic dataset from Zymo Microbial Community Standard. Medaka was run after Racon and Homopolish was invoked after Medaka.

## Discussion

This paper proposed a Nanopore-specific polishing program called Homopolish based on a machine learning model trained from homologous sequences. While most state-of-the-art polishing polishing models are trained from Nanopore reads or signals (e.g., Nanopolish, Medaka, HELEN), we showed that homologs conserved in closely-related species provide valuable features for correction of Nanopore systematic errors (e.g., frameshift is extremely rare within coding regions). In addition, because Nanopore signals may be disturbed by species-specific DNA modifications, construction of an universal model for polishing all species is challenging. This work suggested a better model aware of species-specific errors is possible, if both reads and homologous sequences can be incorporated into the underlying training framework (e.g., RNN in Medaka).

In terms of speed, because existing polishing programs (i.e., Medaka and HELEN) were based on deep neural network, GPU acceleration is often required. On the other hand, CPU is sufficient for Homopolish which is based on a traditional support vector machine (e.g., ∼five minutes for polishing a bacterial genome). Although deep neural network is theoretically suitable for learning non-trivial features, we provided a set of manually-inspected features capable of capturing Nanopore systematic errors that may be directly used by other model developer.

Homopolish has been extensively tested over a large number of bacteria and a limited set of virus and fungi. However, we note that the correction power might be reduced on plasmids shaped by mobile genetic elements (e.g., integrons), which exhibit little or no structural conservation across plasmids. Second, because the precompiled RefSeq sketches are from nearly complete genomes, when polishing highly fragmented genomes (e.g., due to low sequencing quality or coverage), the species identification stage (i.e., Minhash) becomes less reliable. Consequently, the user should specify the genus or specie name when using Homopolish for correcting highly-fragmented assembly.

The results indicated accuracy of R10.3 is indeed better than R9.4 on the same metagenomic dataset (Supplementary Tables S7 and S9). We note that, because of using an old version of Guppy (v2.2) basecaller in the metagenomic dataset, the read accuracy is stil low (∼Q9-Q10). However, the accuracy of Nanopore reads has been significantly improved recently (e.g., ∼Q13-14 on Guppy v3.6). the genome quality based on latest basecaller also elevated (results not shown). In terms of the final genome quality, the accuracy gaps between R9.4 and R10.3 is not much (i.e., both can exceeds Q50). As the throughput of R10.3 is currently lower than that of R9.4, R9.4 may be still preferred in sequencing projects pooling large samples.

## Methods

Homopolish can be run with or without specifying specie or genus. The entire workflow of Homopolish is illustrated in Figure 6 and briefly explained below. Without specifying specie or genus, the genome to be polished is screened against the virus, bacteria, or fungus genomes compressed in (MinHash) sketches, which are pre-compiled from NCBI RefSeq database using Mash [15]. If genus or specie name is specified, random *t* genomes (default *t* = 20) of the same specie or genus will be downloaded from NCBI. Subsequently, the retrieved genomes are aligned against the draft genome in order to extract conserved homologous sequences. A number of features were extracted from the alignments and used for training and for predicting the correct bases at each locus.

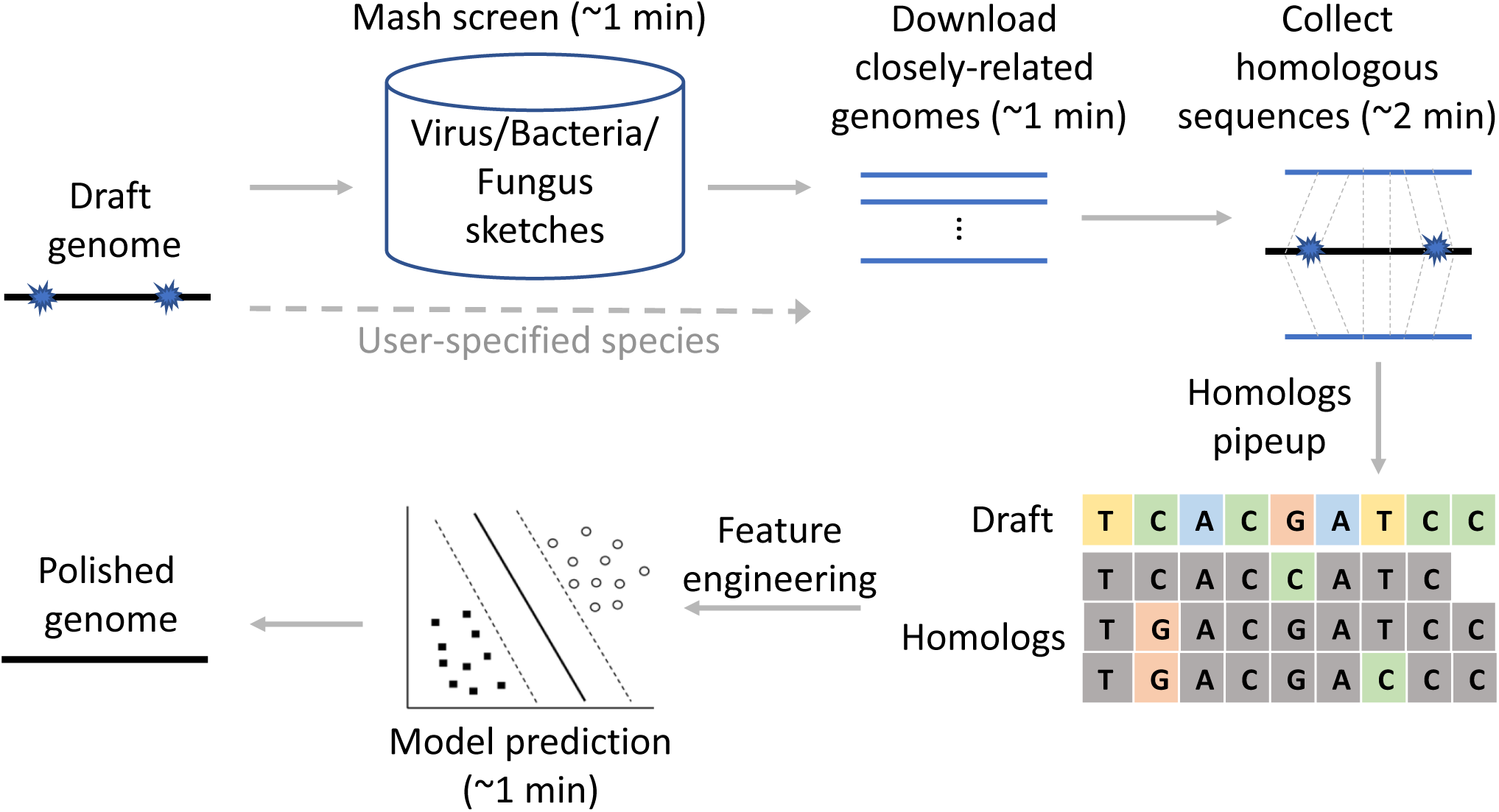

### Retrieval of homologous sequences via MinHash sketches

Given a genome *G* to be polished, Homopolish first identifies, downloads, and extracts homologous sequences from closely-related genomes. In practice, whole-genome alignment against the entire NCBI RefSeq genome database is time-consuming and infeasible. Instead, these genomes are compressed into reduced representation called Mash sketches [15]. Specifically, the genomes were first downloaded from RefSeq and compiled into 1,000 sketches for each genome. The similarity of *G* against all RefSeq genomes can be thus estimated in seconds by Mash. Only the genomes with identity at least *p* (default 95%) are considered to be downloaded. Subsequently, top *t* (default 20) genomic sequences are automatically downloaded from NCBI, which takes around one minute for retrieving 20 bacterial genomes. Alternatively, Homopolish allows the user to specify genus and specie names of genome *G*, and *t* random genomes of the same species or genus will be downloaded from NCBI. Alternatively, random *t* genomes can be directly downloaded without MinHash screening if the genus or specie name is specified by the user. Finally, these downloaded genomes will be aligned to the genome *G* via minimap2 (with options asm5) [16], in order to identify homologous sequences as well as the alignment profile (i.e., pileups) against *G*.

### Feature engineering and model training

A number of features were extracted from the homologous alignment profile in order to distinguish Nanopore systematic errors from inter-strain variations. These features were then used for training and for prediction over a support vector machine (SVM).

### Feature engineering

At each locus of genome *G*, twelve features were extracted from homologous alignment profile for distinguishing Nanopore systematic errors from inter-strain variations. Note that we do not polish mismatched errors as they are frequently occurred in different strains of the same species and are less destructive compared with homopolymer (indels) errors. The first nine features are matched-allele counts of A,T,C,G, inserted-allele counts of A,T,C,G (Supplementary Figure S5(a)), and deletion-allele counts in homologous sequences (Supplementary Figure S5(b)). These nine features aim to reflect the degree of conservation measured by allele frequencies within homologs (e.g., a single allele with 100% frequency indicates a very conservative locus). The 10^*th*^ feature is termed homologous coverage (similar to read coverage), which reflects the confidence of homologous allele counts (i.e., higher coverage stands for higher confidence). The above ten features are min-max normalized into [0,1] intervals.

The eleventh feature reflects homopolymer length with respect to the base being predicted. Supplementary Figure S6(a) illustrates an insertion error at the start of consecutive five adenine bases (i.e., length five homopolymer). This feature is important as most Nanopore systematic errors are found around homopolymers. Unexpectedly, we observed one-hot encoding of this feature achieves superior accuracy than naive min-max normalization (see Supplementary Figure S7). We hypothesize this might owing to side effects of basecalling algorithms (e.g., Guppy) overfitting particular lengths of homopolymers [17]. Hence, the length of homopolymer may be better interpreted as categorical instead of numerical feature. In order to limit this feature with fixed dimensions, we observed the homopolymer length is a skewed distribution (Supplementary Figure S6(b)), whereas the majorities are lengths shorter than three and very few exceeds ten. Consequently, the homopolymer length is one-hot encoded into ten categories {1, 2, 3, …, 8, 9, ≥ 10}.

The last feature is called local sequence similarity, which was manually inspected by Integrated Genomics Viewer [18]. We occasionally found that the minor (instead of the major) allele among the homologous sequences are the true allele at several loci, which violate the assumption of first ten allele-count features and were wrongly corrected as the major one. Further investigation indicated that the homologous sequences flanking these minor alleles are more identical to the ground-truth genome (see Supplementary Figure S8). For example, the homologous sequence 1 in Supplementary Figure S8(a) is perfectly identical to the ground-truth genome, and sequences 1 and 4 in Supplementary Figure S8(b) are the same. This is also explicitly or implicitly implemented in SNP calling algorithms (e.g., haplotype counts in freebayes and CNN in DeepVariant [19, 20].

In reality, because the ground-truth genome is not known in advance, the similarity of flanking sequences can only be estimated by aligning onto the draft genome *G*. But we observed the similarity is often indistinguishable between the major/minor alleles. For instance, in Supplementary Figure S8(b), the homologous sequences flanking the major (e.g., 2 and 3) and minor (e.g., 1 and 4) alleles both differ with the draft genome *G* by one insertion. Thus, their identities against draft genome are completely the same. Interestingly, these erroneous loci were mainly found in pairs and in proximity (e.g., Loci 1 and 2 in Supplementary Figures 8(a)(b)), and their major/minor alleles are mutually-exclusive. Although we cannot explain this phenomena, we found the minor (yet more similar with truth genome) alleles are usually the correct base, whereas the major alleles are not. As a result, the minor alleles at these paired loci is encoded as 2, the major as 1, and the other unpaired bases as 0.

### Training a support vector machine

Seven class labels, insertions of A,T,C,G, no insertion, deletion, and no deletion are assigned to each base being trained or predicted. The class of no insertion is defined for feature vectors containing insertion alleles yet the truth label is no insertion. Instead of grouping the classes of no insertion and of no deletion, we observed the feature vectors of these two classes are quite distant owing to independent insertion and deletion feature space (Supplementary Figure S9). Separating these two classes lead to better prediction accuracy than grouping them into a single class (results not shown).

The class frequencies of both training and testing samples are quite imbalanced across the seven classes, whereas the no deletion class dominates all the others (≈30 millions) (Table **??**). We remove duplicated feature vectors belonging to the no deletion class. The level of imbalance is greatly reduced although the samples of no deletion class is still the majority (∼38 thousands). These feature vectors were trained by a support vector machine (SVM) with the Radial Basis Function (RBF) kernel (*c*=1.0) using five-fold cross validations. During prediction, the genome is chopped into 10kbp segments and all segments can be parallelly polished in a multicore environment.

### Sequencing and assembly of Nanopore and Illumina isolates and metagenomic datasets

Seven bacterial isolates (*Pvulgaris* CCU063, *Pvulgaris* GOKU, *K*.*pneumoniae* SAWA, *E*.*anopheles* SUE, *Salgae* HIDE, *Salgae* VGH117) were sequenced using both Illumina and Nanopore with ∼100-300 coverage (Supplementary Tables S1 and S2). The ground truth genomes of the seven bacteria were constructed by hybrid assembly of Nanopore and Illumina reads via Unicyler [4] (Supplementary Tables S4). The Nanopore reads and ground truth genome of *EcoliK*12 were downloaded from [21]. The Nanopore reads of lambda phage virus were downloaded from Short Read Archieve (SRR12602365). The fungus Nanopore reads of *Saccharomyces cerevisiae* CEN.PK113-7D were downloaded from Short Read Archieve (SRR5989372) and RefSeq (GCA 000269885.1) is used as its ground truth genome.

Two Nanopore metagenomic datasets (R9.4 and R10.3) of ZymoBIOMICS Microbial Community Standard were downloaded from Loman Lab [14] (Supplementary Table S3). The original metagenomic dataset includes eight bacteria and two fungi. Lactobacillus was excluded because of unusually-low quality (∼Q20) possibly due to inconsistent strains. The two fungi were removed owing to highly fragmented assembly.

The Nanopore reads of bacterial isolates, virus, and fungi were first trimmed by PoreChop and assembled into chromosomes by Flye (v2.7) [7] (Supplementary Table S6), whereas plasmids are excluded in subsequent analysis. The metagenomic dataset were assembled by MetaFlye (v2.7) [12] (Supplementary Table S4). All these assembled genomes were first polished by either Racon or MarginPolish for correcting simple sequencing errors. The remaining systematic errors were then removed by either Medaka, HELEN, or Homopolih. The quality of polishing genome (Q score) as well as numbers of insertions, deletions, and mismatches were calculated by fastmer [13].

## Program availability

Homopolish is freely available at https://github.com/ythuang0522/homopolish

## Author’s contributions

YTH design the study. PWS and IPC implemented the python. PYL provided bacterial sequencing. YTH wrote the masnucript.

