## Supplementary Material for "High-Quality Genomes of Nanopore Sequencing by Homologous Polishing"

### Supplementary Material for High-Quality Genome from Nanopore Sequencing by Homologous Polishing

September 20, 2020

### 1 Supplementary Figures

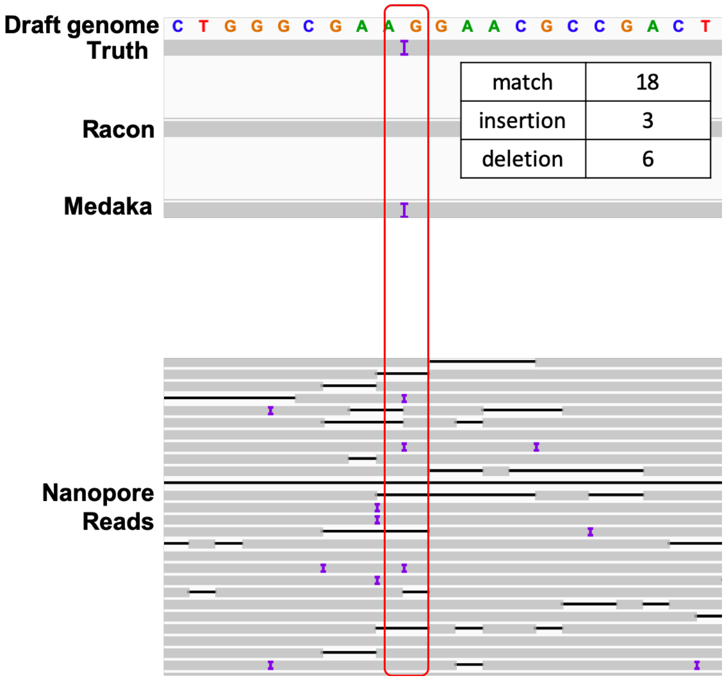


Figure S1: Illustration of Nanopore systematic errors after polishing by Racon and by Medaka. The top shows the genomes before and after polishing. The bottom shows the IGV read alignments. Because majority of reads suggested no insertions at this locus (i.e., 18 vs 3 reads), Racon's majority-based strategy is unable to fix this error. Medaka successfully correct this systematic error by adding the missed nucleotide.

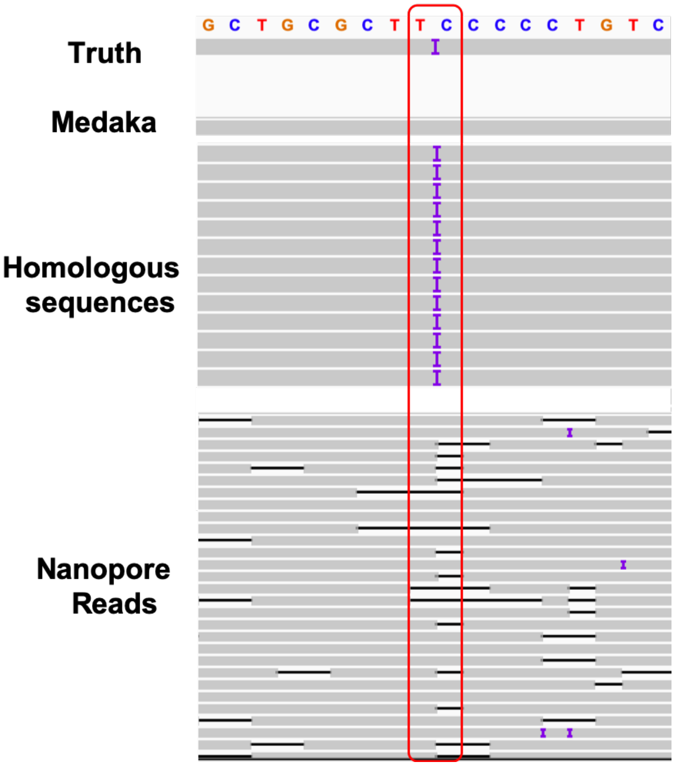


Figure S2: Illustrations of Nanopore systematic errors left by Medaka polishing and importance of homologous sequences. Because all the reads indicate no insertion at this locus (see bottom), Medaka failed to correct this sort of systematic errors. On the other hand, the homologous sequences at this region all agreed on an insertion at this locus, which in turn fix this systematic error.

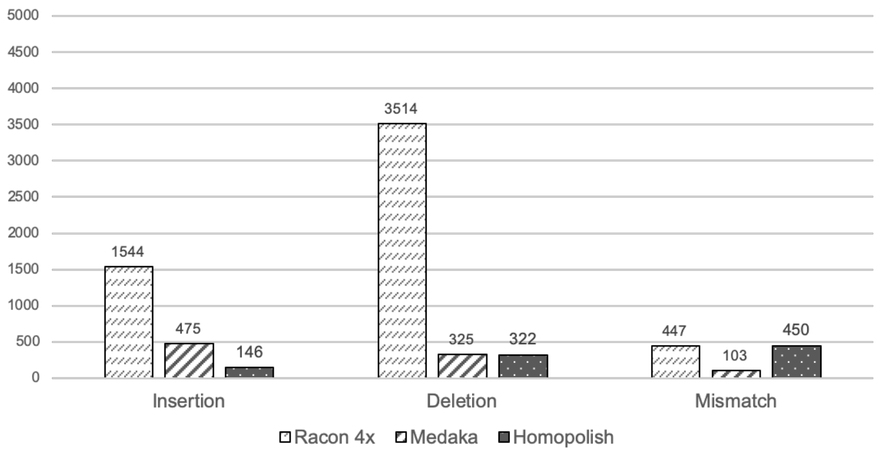


Figure S3: Numbers of indels and mismatches of Bacillus genome after polishing by Racon, by Medaka, and by Homopolish.

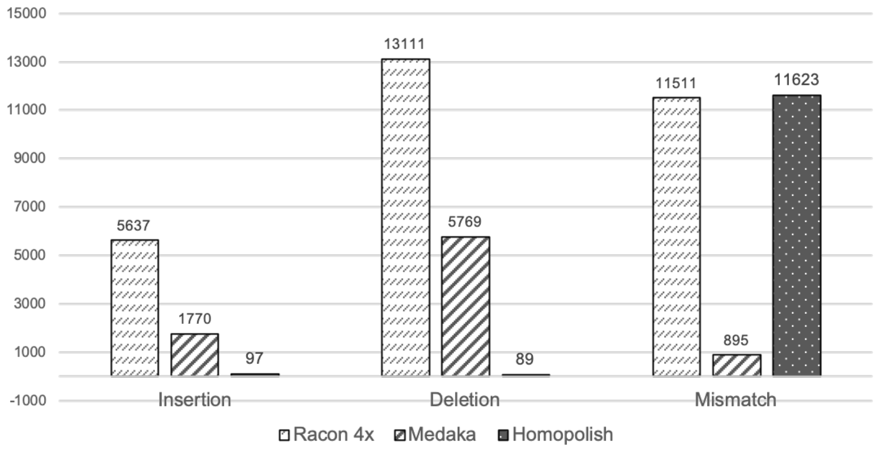


Figure S4: Numbers of indels and mismatches of klebsiella pneumonia SAWA genome after polishing by Racon, by Medaka, and by Homopolish.

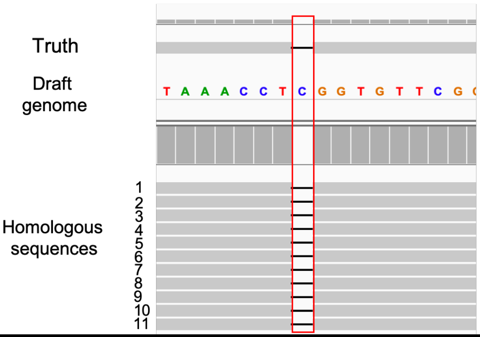


| A | T | C | G | gap | Ins_A | Ins_T | Ins_C | Ins_G | Coverage | Label |
|---|---|---|---|---|---|---|---|---|---|---|
| 0 | 0 | 0 | 0 | 11 | 0 | 0 | 0 | 0 | 11 | Deletion |

(a) Example of a deletion alignment profile and associated feature vector

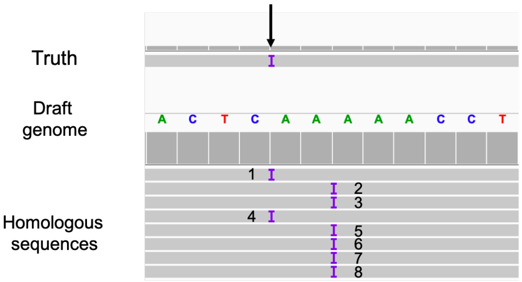


| A | T | C | G | gap | Ins_A | Ins_T | Ins_C | Ins_G | Coverage | Label |
|---|---|---|---|---|---|---|---|---|---|---|
| 0 | 0 | 0 | 0 | 0 | 2 | 0 | 0 | 0 | 8 | Insertion A |

(b) Example of an insertion alignment profile and feature vector

Figure S5: Example of feature vectors for insertions and deletions. The allele counts of each feature are shown in the corresponding count tables.

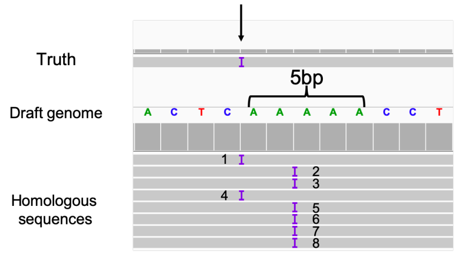


| A | T | C | G | gap | Ins_A | Ins_T | Ins_C | Ins_G | Coverage | Homopolymer length |
|---|---|---|---|---|---|---|---|---|---|---|
| 0 | 0 | 0 | 0 | 0 | 2 | 0 | 0 | 0 | 8 | 5 |

(a) A systematic error surrounding a homopolymer

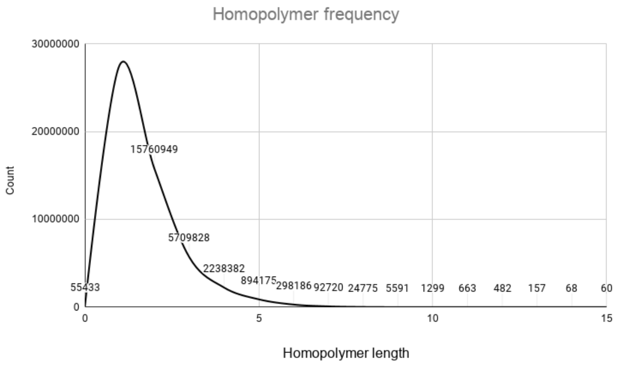


(b) Frequency distribution of homologous lengths

Figure S6: Illustration of Nanopore systematic errors surrounding homopolymers. (a) An insertion error at the start of a homopolymer run of five adenine bases; (b) the frequency distribution of homopolymer lengths in the genome.

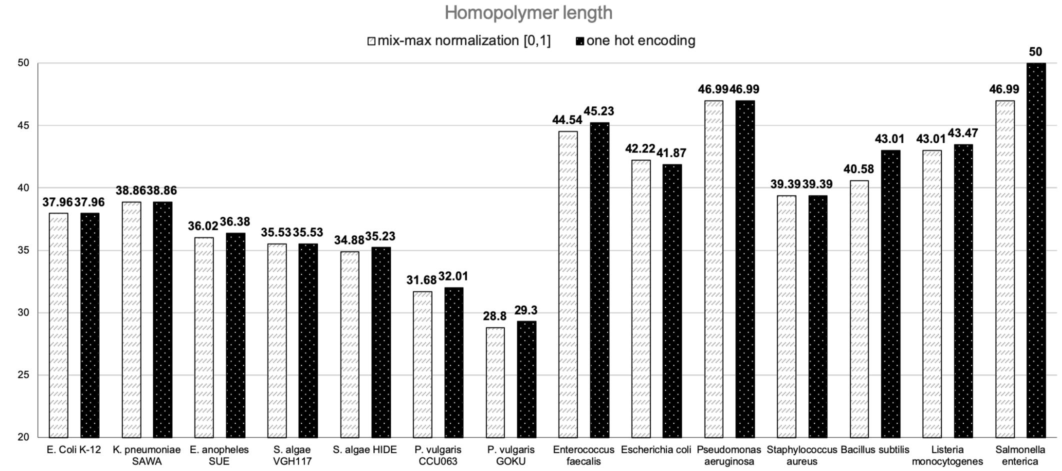


Figure S7: Accuracy comparison of min-max normalization with one-hot encoding of the homopolymer length feature.

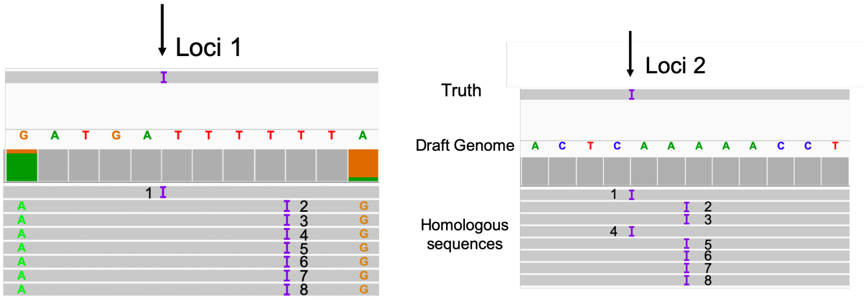


(a) Distinct sequence similarity flanking the major/minor alleles

(b) Identical sequence similarity flanking the major/minor alleles

Figure S8: Illustration of sequence similarity flanking the major/minor alleles. (a) An example of distinct similarity. The homologous sequence 1 flanking the minor allele is more similar to the draft genome, while those flanking the major allele contain mismatches and insertions. (b) An example of identical similarity. The homologous sequences flanking the major (2,3,5-8) and the minor (1,4) contain one insertion to the draft genome.

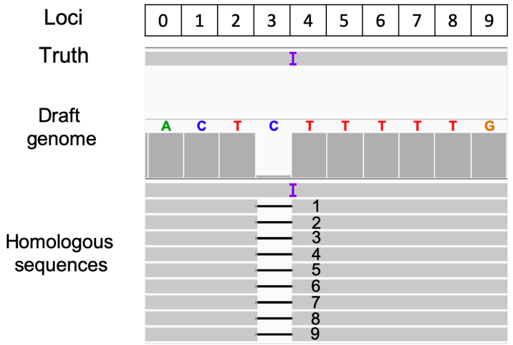


Feature Vectors

| Loci | A | T | C | G | gap | Ins_A | Ins_T | Ins_C | Ins_G | Coverage | Label |
|---|---|---|---|---|---|---|---|---|---|---|---|
| 3 | 0 | 0 | 0 | 0 | 9 | 0 | 0 | 0 | 0 | 10 | No Correction |
| 3 | 0 | 0 | 0 | 0 | 0 | 0 | 1 | 0 | 0 | 10 | T |

Figure S9: Comparison of feature vectors of a no deletion and a no feature vector. Although truth genome indicates no need of correction at both loci, the feature vector containing the deletion alleles is unrelated to that containing the insertion.

### 2 Supplementary Tables

This selection lists the supplementary tables of sequencing, assembly, and polished results of each program on metagenomic or isolate datasets.

Table S1: In-house Nanopore isolate sequencing statistic of six bacteria. The read numbers, N50 read length, maximum read length, and sum of total read bases are listed for each strain.

<table>
<tr><th colspan="5">Nanopore sequencing</th></tr>
<tr><th>strain</th><th>Num.</th><th>N50</th><th>Max</th><th>Sum</th></tr>
<tr><td>K. pneumoniae SAWA</td><td rowspan="3">406,263</td><td rowspan="3">17,529</td><td rowspan="3">122,092</td><td rowspan="3">842,252,640</td></tr>
<tr><td>E. anopheles SUE</td></tr>
<tr><td>S. algae VGH117</td></tr>
<tr><td>S. algae HIDE</td><td>178,071</td><td>25,496</td><td>141,550</td><td>2,944,017,556</td></tr>
<tr><td>P. vulgaris CCU063</td><td>161,898</td><td>10,271</td><td>49,577</td><td>1,292,396,754</td></tr>
<tr><td>P. vulgaris GOKU</td><td>33,172</td><td>9,292</td><td>79,283</td><td>163,089,117</td></tr>
</table>

Table S2: In-house Illumina isolate sequencing statistic of six bacteria. The read numbers, N50 read length, maximum read length, and sum of total read bases are listed for each strain.

| Illumina sequencing | | | | |
|---|---|---|---|---|
| strain | Num. | N50 | Max | Sum |
| K. pneumoniae SAWA | 2,204,942 | 151 | 151 | 332,946,242 |
| E. anopheles SUE | 2,518,328 | 151 | 151 | 380,267,528 |
| S. algae VGH117 | 2,295,502 | 151 | 151 | 346,620,802 |
| S. algae HIDE | 2,166,486 | 151 | 151 | 327,139,386 |
| P. vulgaris CCU063 | 2,647,252 | 151 | 151 | 399,735,052 |
| P. vulgaris GOKU | 2,723,350 | 151 | 151 | 411,225,850 |

Table S3: Public Nanopore metagenomic sequencing statistics from the ZymoBIOMICS Microbial Community Standard sequenced by Loman et al. using R9.4 and R10.4 flowcells. The read numbers, N50 read length, maximum read length, and sum of total read bases are listed for each strain.

| Version | Num. | N50 | Max | Sum |
|---|---|---|---|---|
| R9.4 | 3,238,505 | 5,331 | 320,098 | 13.63G |
| R10.3 | 1,160,526 | 26,245 | 249,289 | 4.4G |

Table S4: Statistic of reference genomes of in-house sequenced isolates (ground truth) reconstructed via hybrid assembly. The contig numbers and genome sizes are listed for each strain.

| Isolate dataset ground truth | | |
|---|---|---|
| strain | contig num. | genome size |
| K. pneumoniae SAWA | 1 | 5,403,374 |
| E. anopheles SUE | 1 | 4,201,198 |
| S. algae VGH117 | 1 | 4,796,801 |
| S. algae HIDE | 1 | 4,950,784 |
| P. vulgaris CCU063 | 1 | 4,141,308 |
| P. vulgaris GOKU | 1 | 4,141,320 |

Table S5: Statistics of metagenomic assembly from the ZymoBIOMICS dataset

| Assembly data | | |
|---|---|---|
| strain | n | size |
| Enterococcus faecalis | 1 | 2,867,861 |
| Escherichia coli | 1 | 4,780,323 |
| Pseudomonas aeruginosa | 1 | 6,800,605 |
| Staphylococcus aureus | 1 | 2,737,456 |
| Bacillus subtilis | 1 | 4,069,116 |
| Salmonella enterica | 1 | 4,774,301 |
| Listeria monocytogenes | 1 | 3,013,178 |

Table S6: Assembly statistics of in-house isolates by Nanopore-only sequencing.

| Assembly data | | |
|---|---|---|
| strain | n | size |
| K. pneumoniae SAWA | 1 | 5,419,444 |
| E. anopheles SUE | 1 | 4,214,907 |
| S. algae VGH117 | 1 | 4,809,726 |
| S. algae HIDE | 1 | 4,963,570 |
| P. vulgaris CCU063 | 1 | 4,153,753 |
| P. vulgaris GOKU | 1 | 4,172,433 |
| E. coli K-12 | 1 | 4,637,729 |

Table S7: Comparison of Q scores and numbers of mismatches/indel errors for Flye, Racon, Medaka, MarginPolish, HELEN, and Homopolish over the ZymoBIOMICS Microbial Community Standard metagenomic dataset sequenced by R9.4 flowcells

| Species | Methods | Q score | Median Qscore | Mismatchs | Insertions | Deletions |
|---|---|---|---|---|---|---|
| *Enterococcus faecalis* | Flye | 22.1 | 22.07 | 389 | 17007 | 253 |
| | Racon 4x | 27.12 | 28.51 | 414 | 1303 | 3808 |
| | Racon 4x + Medaka | 37.18 | 37.7 | 131 | 253 | 157 |
| | Racon 4x + Hompolish | 31.74 | 38.24 | 420 | 57 | 1431 |
| | MarginPolish | 33.74 | 33.98 | 69 | 754 | 372 |
| | MarginPolish + HELEN | 41.87 | 43.98 | 34 | 63 | 86 |
| | MarginPolish + Homopolish | 43.47 | 50 | 71 | 33 | 23 |
| | Racon 4x + Medaka + Homopolish | 42.06 | 45.23 | 132 | 25 | 19 |
| | MarginPolish + HELEN + Homopolish | 43.67 | 50 | 53 | 27 | 41 |
| *Escherichia coli* | Flye | 25.23 | 25.46 | 603 | 11115 | 310 |
| | Racon 4x | 30.33 | 31 | 685 | 488 | 2531 |
| | Racon 4x + Medaka | 35.2 | 37.96 | 527 | 422 | 260 |
| | Racon 4x + Hompolish | 36.33 | 39.39 | 697 | 122 | 112 |
| | MarginPolish | 34.31 | 36.02 | 464 | 521 | 494 |
| | MarginPolish + HELEN | 35.63 | 38.54 | 477 | 229 | 384 |
| | MarginPolish + Homopolish | 37.48 | 41.55 | 484 | 132 | 97 |
| | Racon 4x + Medaka + Homopolish | 37.24 | 41.87 | 521 | 130 | 104 |
| | MarginPolish + HELEN + Homopolish | 37.44 | 41.25 | 471 | 127 | 120 |
| *Pseudomonas aeruginosa* | Flye | 26.39 | 26.5 | 738 | 14394 | 439 |
| | Racon 4x | 32.78 | 33.19 | 622 | 538 | 2417 |
| | Racon 4x + Medaka | 36.54 | 37.96 | 476 | 704 | 324 |
| | Racon 4x + Hompolish | 39.43 | 43.01 | 622 | 24 | 128 |
| | MarginPolish | 34.93 | 35.85 | 449 | 1096 | 637 |
| | MarginPolish + HELEN | 41.48 | 46.99 | 149 | 78 | 256 |
| | MarginPolish + Homopolish | 40.74 | 50 | 440 | 7 | 126 |
| | Racon 4x + Medaka + Homopolish | 40.55 | 46.99 | 474 | 4 | 120 |
| | MarginPolish + HELEN + Homopolish | 42.81 | 90 | 205 | 5 | 146 |
| *Staphylococcus aureus* | Flye | 22.88 | 22.9 | 876 | 12911 | 177 |
| | Racon 4x | 29.96 | 29.94 | 365 | 1012 | 1341 |
| | Racon 4x + Medaka | 35.79 | 36.2 | 247 | 274 | 190 |
| | Racon 4x + Hompolish | 37.31 | 37.83 | 367 | 38 | 96 |
| | MarginPolish | 33.07 | 33.37 | 232 | 845 | 253 |
| | MarginPolish + HELEN | 38.1 | 38.7 | 61 | 172 | 184 |
| | MarginPolish + Homopolish | 38.52 | 40 | 232 | 23 | 124 |
| | Racon 4x + Medaka + Homopolish | 38.69 | 39.39 | 247 | 24 | 93 |
| | MarginPolish + HELEN + Homopolish | 41.25 | 44.58 | 63 | 21 | 118 |
| *Bacillus subtilis* | Flye | 23.31 | 23.31 | 347 | 18254 | 347 |
| | Racon 4x | 28.66 | 28.64 | 447 | 1544 | 3514 |
| | Racon 4x + Medaka | 36.51 | 37.21 | 103 | 475 | 325 |
| | Racon 4x + Hompolish | 36.44 | 36.78 | 450 | 146 | 322 |
| | MarginPolish | 34.77 | 35.23 | 36 | 677 | 635 |
| | MarginPolish + HELEN | 37.28 | 37.7 | 30 | 317 | 409 |
| | MarginPolish + Homopolish | 40.6 | 45.23 | 49 | 102 | 201 |
| | Racon 4x + Medaka + Homopolish | 40.17 | 43.01 | 103 | 95 | 191 |
| | MarginPolish + HELEN + Homopolish | 40.83 | 43.98 | 31 | 89 | 214 |
| *Salmonella enterica* | Flye | 25.23 | 25.25 | 203 | 11446 | 280 |
| | Racon 4x | 30.72 | 31.08 | 339 | 448 | 2611 |
| | Racon 4x + Medaka | 37.3 | 38.86 | 190 | 340 | 217 |
| | Racon 4x + Hompolish | 40.18 | 43.01 | 340 | 14 | 31 |
| | MarginPolish | 34.81 | 35.53 | 63 | 637 | 623 |
| | MarginPolish + HELEN | 37.77 | 38.54 | 36 | 214 | 417 |
| | MarginPolish + Homopolish | 45.31 | 50 | 70 | 20 | 28 |
| | Racon 4x + Medaka + Homopolish | 42.51 | 50 | 186 | 13 | 26 |
| | MarginPolish + HELEN + Homopolish | 46.72 | 50 | 36 | 13 | 36 |
| *Listeria monocytogenes* | Flye | 22.65 | 22.58 | 174 | 11034 | 120 |
| | Racon 4x | 27.5 | 28.81 | 267 | 1277 | 2153 |
| | Racon 4x + Medaka | 31.82 | 38.54 | 94 | 669 | 605 |
| | Racon 4x + Hompolish | 31.87 | 38.54 | 266 | 540 | 545 |
| | MarginPolish | 31.24 | 36.2 | 42 | 846 | 675 |
| | MarginPolish + HELEN | 32.03 | 39.68 | 44 | 655 | 603 |
| | MarginPolish + Homopolish | 32.68 | 46.99 | 42 | 538 | 540 |
| | Racon 4x + Medaka + Homopolish | 32.48 | 43.47 | 94 | 538 | 541 |
| | MarginPolish + HELEN + Homopolish | 32.67 | 46.99 | 44 | 539 | 540 |

Table S8: Comparison of Q scores and numbers of mismatches/indel errors for Flye, Racon, Medaka, MarginPolish, HELEN, and Homopolish on bacterial isolates datasets sequenced by R9.4 flowcells

| Species | Methods | Avg Q score | Median Q score | Mismatchs | Insertions | Deletions |
|---|---|---|---|---|---|---|
| *E. Coli K-12* | Flye | 22.09 | 22.25 | 14241 | 6248 | 8193 |
| | Racon 4x | 21.68 | 21.83 | 10743 | 3349 | 17450 |
| | Racon 4x + Medaka | 25.85 | 26.22 | 708 | 830 | 10537 |
| | Racon 4x + Hompolish | 25.92 | 26.31 | 10742 | 201 | 917 |
| | MarginPolish | 21.94 | 22.11 | 10742 | 2076 | 16893 |
| | MarginPolish + HELEN | 20.77 | 20.73 | 8918 | 4200 | 21383 |
| | MarginPolish + Homopolish | 25.92 | 26.35 | 10738 | 191 | 941 |
| | Racon 4x + Medaka + Homopolish | 34.15 | 37.96 | 708 | 181 | 894 |
| | MarginPolish + HELEN + Homopolish | 26.18 | 26.19 | 9210 | 290 | 716 |
| *K. pneumoniae SAWA* | Flye | 20.79 | 20.78 | 16431 | 22205 | 6255 |
| | Racon 4x | 22.49 | 22.49 | 11511 | 5637 | 13111 |
| | Racon 4x + Medaka | 28.06 | 28.1 | 895 | 1770 | 5769 |
| | Racon 4x + Hompolish | 26.6 | 26.52 | 11623 | 97 | 89 |
| | MarginPolish | 22.8 | 22.8 | 9778 | 3405 | 14925 |
| | MarginPolish + HELEN | 22.27 | 22.22 | 7904 | 4776 | 19117 |
| | MarginPolish + Homopolish | 27.31 | 27.33 | 9874 | 64 | 88 |
| | Racon 4x + Medaka + Homopolish | 37.34 | 38.86 | 896 | 57 | 43 |
| | MarginPolish + HELEN + Homopolish | 28.14 | 28.18 | 7983 | 74 | 232 |
| *E. anopheles SUE* | Flye | 22.25 | 22.2 | 1363 | 18718 | 5039 |
| | Racon 4x | 23.57 | 23.57 | 1163 | 6097 | 11215 |
| | Racon 4x + Medaka | 26.43 | 26.48 | 566 | 1114 | 7891 |
| | Racon 4x + Hompolish | 33.11 | 33.57 | 1224 | 384 | 443 |
| | MarginPolish | 25.57 | 25.61 | 389 | 1055 | 10198 |
| | MarginPolish + HELEN | 23.89 | 23.91 | 669 | 4614 | 11753 |
| | MarginPolish + Homopolish | 35.8 | 37.21 | 447 | 222 | 435 |
| | Racon 4x + Medaka + Homopolish | 35.32 | 36.38 | 611 | 216 | 407 |
| | MarginPolish + HELEN + Homopolish | 33.57 | 34.44 | 723 | 342 | 774 |
| *S. algae VGH117* | Flye | 23.74 | 23.7 | 1307 | 15982 | 3054 |
| | Racon 4x | 26.48 | 26.53 | 1262 | 3103 | 6423 |
| | Racon 4x + Medaka | 29.58 | 29.69 | 558 | 1028 | 3695 |
| | Racon 4x + Hompolish | 32.3 | 33.01 | 1399 | 562 | 863 |
| | MarginPolish | 28.08 | 28.11 | 582 | 989 | 5896 |
| | MarginPolish + HELEN | 27.6 | 27.66 | 523 | 1370 | 6449 |
| | MarginPolish + Homopolish | 33.76 | 34.69 | 702 | 407 | 907 |
| | Racon 4x + Medaka + Homopolish | 34.57 | 35.53 | 639 | 340 | 696 |
| | MarginPolish + HELEN + Homopolish | 33.59 | 35.02 | 659 | 424 | 1014 |
| *S. algae HIDE* | Flye | 23.97 | 23.92 | 1122 | 15674 | 2937 |
| | Racon 4x | 26.16 | 26.16 | 1034 | 2459 | 8414 |
| | Racon 4x + Medaka | 29.7 | 29.79 | 465 | 1013 | 3790 |
| | Racon 4x + Hompolish | 32.42 | 33.23 | 1189 | 508 | 1117 |
| | MarginPolish | 28.24 | 28.23 | 438 | 854 | 6088 |
| | MarginPolish + HELEN | 27.87 | 27.95 | 391 | 1187 | 6453 |
| | MarginPolish + Homopolish | 34.48 | 35.02 | 571 | 284 | 897 |
| | Racon 4x + Medaka + Homopolish | 35.05 | 35.23 | 541 | 283 | 714 |
| | MarginPolish + HELEN + Homopolish | 33.91 | 34.26 | 552 | 369 | 1076 |
| *P. vulgaris CCU063* | Flye | 23.5 | 23.56 | 783 | 14945 | 2819 |
| | Racon 4x | 24.21 | 24.2 | 1047 | 2767 | 11904 |
| | Racon 4x + Medaka | 27.68 | 27.75 | 345 | 458 | 6266 |
| | Racon 4x + Hompolish | 28.72 | 29.05 | 1716 | 1228 | 2620 |
| | MarginPolish | 26.58 | 26.62 | 170 | 874 | 8064 |
| | MarginPolish + HELEN | 26.58 | 26.68 | 169 | 1283 | 7663 |
| | MarginPolish + Homopolish | 31.18 | 31.46 | 635 | 662 | 1861 |
| | Racon 4x + Medaka + Homopolish | 31.51 | 32.01 | 710 | 567 | 1647 |
| | MarginPolish + HELEN + Homopolish | 30.71 | 31.02 | 711 | 767 | 2040 |
| *P. vulgaris GOKU* | Flye | 20.56 | 20.5 | 1501 | 33165 | 2059 |
| | Racon 4x | 20.91 | 20.85 | 3337 | 22954 | 7497 |
| | Racon 4x + Medaka | 23.97 | 24.02 | 1315 | 12180 | 3136 |
| | Racon 4x + Hompolish | 25.85 | 26.5 | 4087 | 4745 | 1939 |
| | MarginPolish | 22.49 | 22.53 | 1127 | 17146 | 5168 |
| | MarginPolish + HELEN | 26.58 | 26.68 | 169 | 1283 | 7663 |
| | MarginPolish + Homopolish | 27.77 | 28.39 | 1846 | 3685 | 1397 |
| | Racon 4x + Medaka + Homopolish | 28.77 | 29.3 | 1830 | 2572 | 1096 |
| | MarginPolish + HELEN + Homopolish | 24.32 | 24.84 | 4979 | 4526 | 4887 |

Table S9: Comparison of numbers of mismatch, insertion, and deletion errors produced by Flye, Racn, Medka, and Homopolish on one metagenomic dataset of seven species and one isolate (E coli K12 MG1655) sequenced by R10.3 flowcells

| Species | Methods | Avg. Q score | Median Q score | Mismatchs | Insertions | Deletions |
|---|---|---|---|---|---|---|
| Enterococcus faecalis | Flye | 39.45 | 40 | 70 | 59 | 194 |
| | Racon 4x | 39.52 | 40.46 | 94 | 116 | 108 |
| | Racon 4x + Medaka | 44.54 | 50 | 57 | 21 | 22 |
| | Racon 4x + Medaka + Homopolish | 45.2 | 90 | 57 | 23 | 6 |
| Escherichia coli | Flye | 34.45 | 35.45 | 749 | 175 | 659 |
| | Racon 4x | 34.46 | 35.69 | 920 | 237 | 533 |
| | Racon 4x + Medaka | 37.86 | 41.55 | 559 | 84 | 129 |
| | Racon 4x + Medaka + Homopolish | 38.15 | 40.97 | 547 | 92 | 83 |
| Pseudomonas aeruginosa | Flye | 36.57 | 37.96 | 480 | 149 | 864 |
| | Racon 4x | 37.5 | 39.59 | 591 | 104 | 511 |
| | Racon 4x + Medaka | 40.46 | 50 | 420 | 48 | 142 |
| | Racon 4x + Medaka + Homopolish | 41.01 | 90 | 420 | 0 | 118 |
| Staphylococcus aureus | Flye | 28.39 | 28.57 | 999 | 2750 | 195 |
| | Racon 4x | 27.43 | 27.43 | 1474 | 3020 | 428 |
| | Racon 4x + Medaka | 29.46 | 29.38 | 1112 | 1595 | 371 |
| | Racon 4x + Medaka + Homopolish | 33.42 | 33.77 | 1118 | 12 | 106 |
| Bacillus subtilis | Flye | 38.09 | 39.21 | 25 | 92 | 506 |
| | Racon 4x | 38.34 | 40 | 69 | 162 | 362 |
| | Racon 4x + Medaka | 41.68 | 46.99 | 20 | 86 | 169 |
| | Racon 4x + Medaka + Homopolish | 41.97 | 50 | 20 | 80 | 157 |
| Salmonella enterica | Flye | 34.24 | 34.44 | 456 | 623 | 708 |
| | Racon 4x | 35.73 | 35.69 | 380 | 403 | 489 |
| | Racon 4x + Medaka | 46.73 | 50 | 30 | 34 | 37 |
| | Racon 4x + Medaka + Homopolish | 48.32 | 53.01 | 30 | 23 | 17 |
| Listeria monocytogenes | Flye | 32.91 | 42.22 | 54 | 535 | 942 |
| | Racon 4x | 32.72 | 40.54 | 87 | 604 | 909 |
| | Racon 4x + Medaka | 33.32 | 50 | 45 | 527 | 820 |
| | Racon 4x + Medaka + Homopolish | 33.26 | 48.24 | 46 | 535 | 833 |
| E coli K12 (isolate) | Racon 4x + Medaka | 35.97 | 46.99 | 23 | 221 | 928 |
| | Racon 4x + Medaka + Homopolish | 36.28 | 90 | 23 | 182 | 887 |

Table S10: Label frequency before and after removing duplicate feature vectors

| | Frequency | |
|---|---|---|
| Label | before | after |
| Insertion A | 4,683 | 4,683 |
| Insertion T | 4,463 | 4,463 |
| Insertion C | 1,799 | 1,799 |
| Insertion G | 1,625 | 1,625 |
| Deletion | 14,935 | 14,935 |
| No Deletion | 30,208,536 | 38,020 |
| No Insertion | 7,849 | 7,849 |
